# The global and local distribution of RNA structure throughout the SARS-CoV-2 genome

**DOI:** 10.1101/2020.07.06.190660

**Authors:** Rafael de Cesaris Araujo Tavares, Gandhar Mahadeshwar, Anna Marie Pyle

## Abstract

SARS-CoV-2 is the causative viral agent of COVID-19, the disease at the center of the current global pandemic. While knowledge of highly structured regions is integral for mechanistic insights into the viral infection cycle, very little is known about the location and folding stability of functional elements within the massive, ~30kb SARS-CoV-2 RNA genome. In this study, we analyze the folding stability of this RNA genome relative to the structural landscape of other well-known viral RNAs. We present an in-silico pipeline to locate regions of high base pair content across this long genome and also identify well-defined RNA structures, a method that allows for direct comparisons of RNA structural complexity within the several domains in SARS-CoV-2 genome. We report that the SARS-CoV-2 genomic propensity to stable RNA folding is exceptional among RNA viruses, superseding even that of HCV, one of the most highly structured viral RNAs in nature. Furthermore, our analysis reveals varying levels of RNA structure across genomic functional regions, with accessory and structural ORFs containing the highest structural density in the viral genome. Finally, we take a step further to examine how individual RNA structures formed by these ORFs are affected by the differences in genomic and subgenomic contexts. The conclusions reported in this study provide a foundation for structure-function hypotheses in SARS-CoV-2 biology, and in turn, may guide the 3D structural characterization of potential RNA drug targets for COVID-19 therapeutics.

## Introduction

SARS-CoV-2 is an enveloped positive-strand RNA virus and the etiological agent of Covid-19 (Wu et al. 2020), a highly infectious human disease at the center of a world pandemic that has taken thousands of lives worldwide (Casanova et al. 2020; Gates 2020; Sempowski et al. 2020). This virus is a member of the family of coronaviruses, known for having the largest genomes among all RNA viruses (Fehr and Perlman 2015). Almost 30 kb in length (Kim et al. 2020), SARS-CoV-2 RNA genome imposes new challenges to RNA structural biology due to its size and complexity.

Following viral entry and uncoating, the genomic RNA serves as the template for translation of a multicomponent replicase-transcriptase complex that is responsible for synthesizing the viral transcriptome, which includes a series of subgenomic RNAs from which other virion components and accessory protein factors are expressed (Fehr and Perlman 2015). Consistent with reports on other coronaviruses, the SARS-CoV-2 genome contains highly conserved RNA structural elements that likely play pivotal roles in viral replication, including several structures in the UTRs and a ribosomal frameshifting element (Rangan et al. 2020). Although many of these motifs have been functionally studied and extensively modeled in other betacoronaviruses (Chen and Olsthoorn 2010; Yang and Leibowitz 2015; Yang et al. 2015; Madhugiri et al. 2018), very little is known about functional structural elements in the overwhelming majority of the SARS-CoV-2 genome.

In line with previous reports on other coronaviral genomes (Simmonds et al. 2004), SARS-CoV-2 was recently suggested to form genome-scale ordered RNA structure, or GORS (Andrews et al. 2020). As shown in foundational work comparing several families of RNA viruses (Simmonds et al. 2004) and further explored in later studies (Davis et al. 2008; McFadden et al. 2013; Witteveldt et al. 2014), the existence of GORS in positive-strand RNA viruses correlates with features like fitness and persistence. These studies have also established hepaciviral genomes as textbook examples of globally structured RNAs and the most studied member of this genus, hepatitis C virus (HCV), is among the most highly structured viral RNAs known to date. The abundant RNA structures found throughout (mostly) coding regions of that genome not only play individual functional roles (McMullan et al. 2007; Fricke et al. 2015; Pirakitikulr et al. 2016) but also contribute to its higher-order compaction (Davis et al. 2008).

Given the pervasive importance of RNA structure in the various aspects of viral life cycles, it becomes vital to assess the RNA folding stability of SARS-CoV-2 genome relative to other well-known viral RNA systems in order to investigate the biological significance of its RNA structural landscape. Despite recent efforts to identify locally structured regions within the SARS-CoV-2 genome (Andrews et al. 2020; Rangan et al. 2020; Vandelli et al. 2020), the size of this RNA has made it particularly challenging to model its genome-wide secondary structure and, more importantly, to obtain a comparative perspective of the levels of RNA structural complexity across its numerous functional domains. A comprehensive picture of the global and local levels of RNA structure across SARS-CoV-2 genomic transcript would provide a roadmap for comprehending overall structural organization of the virus and would guide identification of regions with the greatest potential to serve as regulatory elements in the virus lifecycle.

In the present work, we show that the potential for stable RNA folding of the SARS-CoV-2 genome supersedes even that of HCV and we discuss the potential biological consequences of this unprecedented level of global structural complexity. We develop a pipeline to extract the base pair content from the modeled secondary structure of any long RNA and to identify regions of well-defined structure across kilobase transcripts. We use this pipeline to scan the SARS-CoV-2 genome and pinpoint the regions of exceptionally significant RNA structural complexity, enabling direct comparisons of predicted structural content among the functional domains of this massive viral genome. We observe a remarkable enrichment of structured regions within ORFs that encode accessory and structural proteins and we elaborate on the potential roles these structures might play in the course of viral infection. Finally, we demonstrate that SARS-CoV-2 ORFs can adopt different structures in the genomic and subgenomic contexts.

We hope that the results and conclusions reported in this study will help design and refine structure-function hypotheses in SARS-CoV-2 biology and that they will guide the identification of RNA drug targets for development of COVID-19 therapeutics.

## Results

### The SARS-CoV-2 genome contains an unprecedented level of stable RNA structure

As an initial global approach to evaluate SARS-CoV-2 RNA structural complexity, we used ScanFold (Andrews et al. 2019) to calculate free-energy Z-scores in windows that were tiled along the entire genome (see Methods) and we analyzed their frequency distribution. In parallel, we performed the same analysis with the HCV genome, which is a hallmark example of globally structured viral RNA and one of the most highly structured RNA genomes ever characterized (Simmonds et al. 2004; Mauger et al. 2015; Pirakitikulr et al. 2016). The West Nile virus was also included for comparison, as viruses in the *Flavivirus* genus are thought to lack globally ordered RNA genomes (Simmonds et al. 2004). Finally, we analyzed a composite set of human mRNAs as a non-viral control believed to lack extensive internal RNA structure (Figure 1).

**Figure 1.**
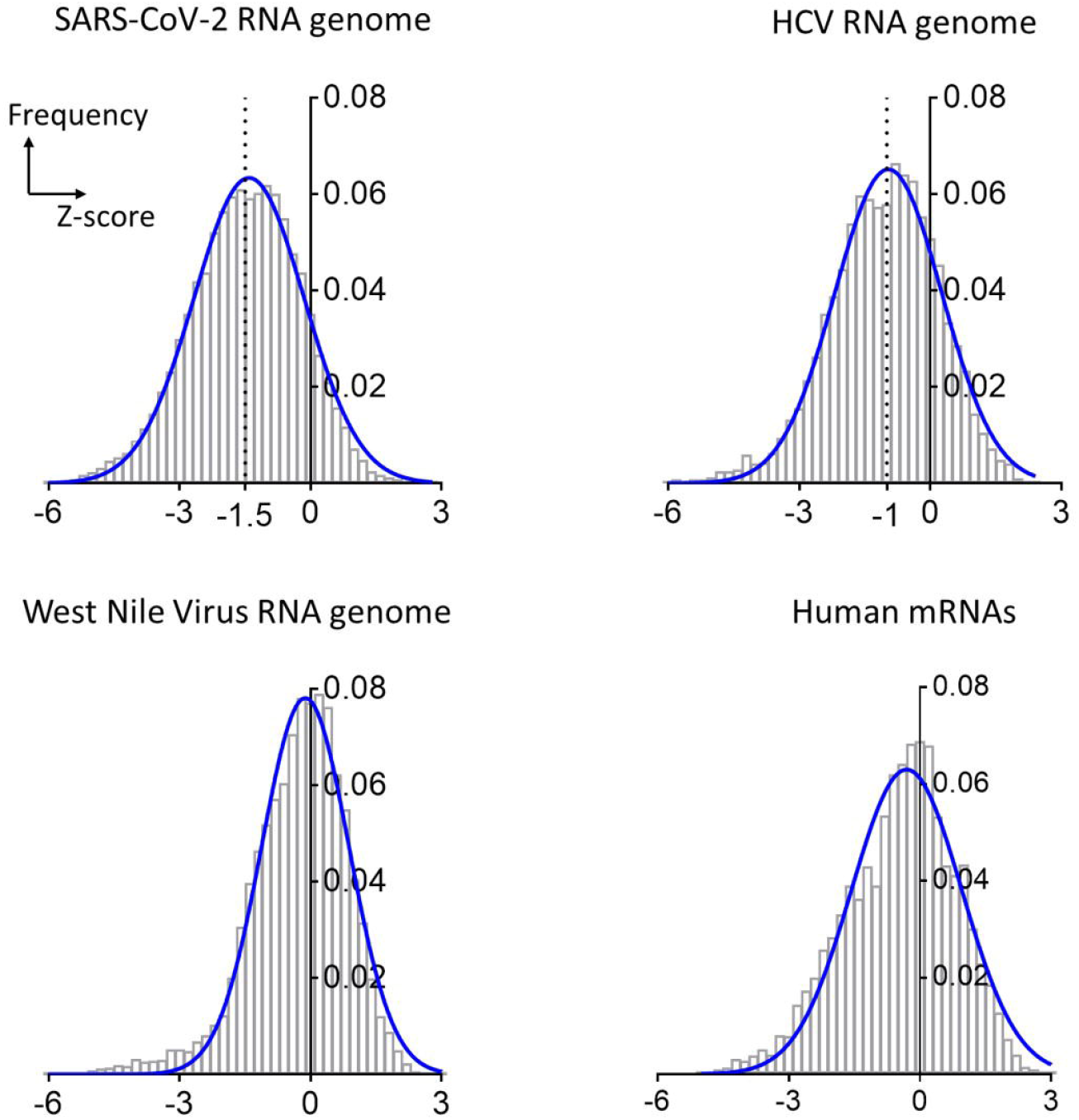
Distributions of Z-scores for the RNA genomes of SARS-CoV-2, HCV and West Nile viruses and a composite of human mRNAs. The bar plots are frequency distributions (y-axis) of free-energy Z-scores (x-axis) calculated in sliding windows tiling each RNA. Each histogram is overlaid with a Gaussian (normal distribution) fit represented by a solid blue curve.

As anticipated, the human mRNAs sampled showed little global tendency to form stable RNA structures (median Z-score −0.35, Figure 1), which is consistent with the presence of local UTR structures and relatively low levels of structure along open reading frames (Mortimer et al. 2014; Wan et al. 2014). In the case of the West Nile virus genome, a Z-score distribution centered at −0.2 (median) similarly suggests the absence of globally ordered RNA folding, in agreement with trends observed for other *Flavivirus* genomes. In very local regions, the human and flaviviral cases genomes displayed a low frequency of highly negative Z-scores (e.g. values below −3, which indicate the presence of highly stable RNA secondary structures), but both distributions suggest the absence of widespread base pairing. By contrast, Z-score distribution for the HCV genome was dominated by negative values (median Z-score −1, Figure 1), indicating a genome-wide propensity to form stable RNA base-pairings. This observation agrees with previous genome-wide analyses of HCV structural content (Simmonds et al. 2004), and studies of discrete RNA secondary structures throughout the HCV UTRs and coding regions (You et al. 2004; McMullan et al. 2007; Friebe and Bartenschlager 2009; Pirakitikulr et al. 2016).

The Z-score distribution for the SARS-CoV-2 genome is shifted far into the negative range (Figure 1), indicating that the genome has a much greater propensity to form stable secondary structures than other RNAs analyzed, by far more than is possible by chance. This is consistent with the reported preference for ordered folding seen in some coronaviral RNAs like MHV (Simmonds et al. 2004). Most strikingly, the SARS-CoV-2 Z-score distribution is centered about a significantly more negative value (median Z-score −1.5) than observed for HCV, suggesting that SARS-CoV-2 genome has almost twice the propensity to form stable base pairings than one of the most structured RNA genomes in Nature, and that it is likely to form extensive secondary structures throughout all of its functional domains, in both coding and noncoding regions. This unusual level of RNA structural stability suggests a vast network of functional RNA structures within the SARS-CoV-2 genome.

### A versatile pipeline for quantifying base pair content within an RNA genome

To map and visualize the entire SARS-CoV-2 RNA structural network, we developed a pipeline for quantitating and comparing relative levels of base pair content and secondary structural features throughout the genome (Figure 2). Initially, we used SuperFold (Smola et al. 2015) to fold the 29.9 kb genome of SARS-CoV-2 in overlapping windows, enabling us to compute a preliminary full-length secondary structure and a genome-wide Shannon entropy profile derived from base pairing probabilities. We then used the resulting secondary structure to calculate the base pair content, or BPC, by scanning the entire RNA in sliding windows (see Methods). We found that the SARS-CoV-2 genome is predominantly folded into stable, discrete secondary structural motifs, with an average BPC of 61% (Figure 2a, median value indicated). This finding agrees with the Z-score analysis which indicated a global propensity for stable structural folding (Figure 1). In order to directly compare the relative structural content of different regions, we also quantified the relative base pair content (BPC_rel_) for each section across SARS-CoV-2 genome (Figure 2b). We define BPC_rel_ as the percentile of BPC at a given site relative to the overall BPC distribution along the length of the RNA (see Methods).

**Figure 2.**
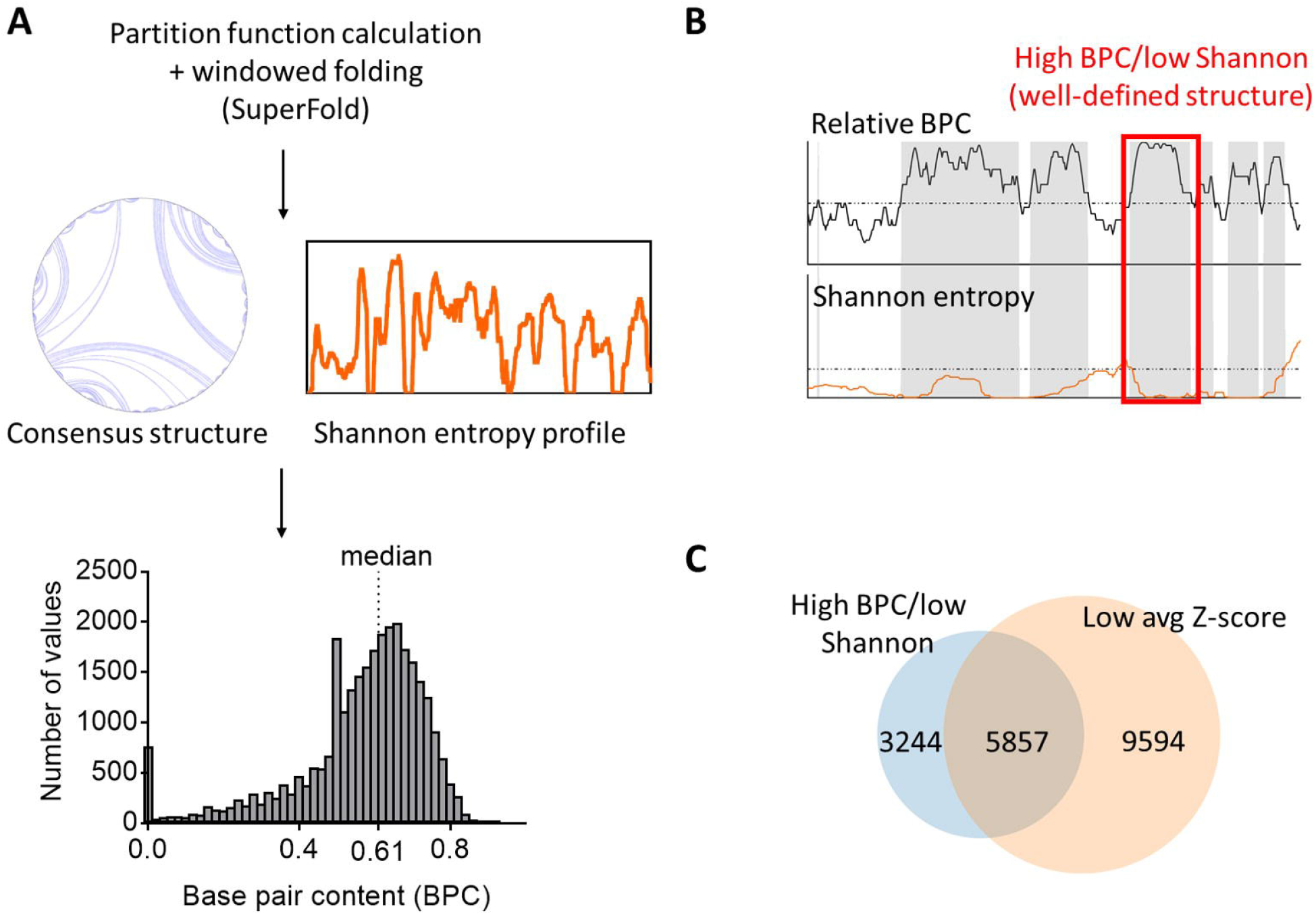
A pipeline to predict and quantify the base pair content across SARS-CoV-2 genome and identify well-defined structured regions. (A) A scheme depicting the steps to predict the secondary structure of SARS-CoV-2 genome in windows using SuperFold. A histogram of base pair content (BPC) values calculated from the predicted secondary structure (gray bar plot) is shown and the median BPC is indicated (0.61). (B) A strategy to identify well-defined structures. The scheme shows shaded regions containing nucleotides that pass two criteria: high relative BPC (upper graph, dashed line indicating the median value of 0.5) and low Shannon entropy (lower graph, dashed line indicating the global Shannon median). The red square highlights one of the regions flagged as forming a well-defined structure. (C) A Venn diagram showing the overlap between the total number of nucleotides identified as having well-defined structure using the procedure in (B) and those nucleotides with low average Z-scores (below the global median) as reported in Andrews et al. (2020).

We then proceeded to sift through the genome, locating discrete regions of well-determined, stable RNA secondary structures. To accomplish this, we adapted a motif discovery method that was originally developed for interpreting the results of chemical probing experiments (Smola et al. 2015). But instead of using SHAPE reactivities as an input, we used the BPC_rel_ distribution across the SARS-CoV-2 genome in conjunction with the corresponding Shannon entropy profile (scheme shown in Figure 2b). This enabled us to flag regions with BPC_rel_ values above 0.5 (i.e. representing BPC values above the predicted global median) and correlate them with Shannon entropy values below the global median, resulting in a metric we define as “high BPC/low Shannon” (see Methods). This definition, which is analogous to the “low SHAPE/low Shannon” designation for flagging probable regions of uniquely determined secondary structure, reveals that 9101 nucleotides, or a third of the entire genome, are located in regions of both high BPC and low Shannon entropy. Since nucleotides with low Shannon entropy are predicted to exist in a single, well-defined folding state (Mathews 2004), high BPC/low Shannon regions are, therefore, clusters of well-defined structures with potential functionality. Our analysis therefore suggests a remarkable abundance of well-defined secondary and tertiary structures within the SARS-CoV-2 genome.

In order to assess the relative thermodynamic stability of specific structured regions defined by this approach, we calculated the relative enrichment in stable base pairs as defined by the ScanFold-Fold analysis in Andrews et al. (Andrews et al. 2020) and computed the overlap between both approaches (Figure 2c). Importantly, we observed that 64% of high BPC/low Shannon entropy regions overlap with regions that have low average Z-scores (also defined here relative to the overall median) and this enrichment is statistically significant (p-value <1e-05, see Methods). In this way, we confirm that the SARS-CoV-2 RNA structural network has a high level of thermodynamic stability.

### The SARS-CoV-2 genome contains specific loci of well-defined RNA structures

Given the abundance of stable secondary structural units that correlate with low Shannon entropy values (high BPC/low Shannon, as defined in Figure 2), we were interested in mapping their distribution across the genome and correlating their location with other units of genomic architecture. To this end, we first calculated the fraction of high BPC/low Shannon nucleotides in bins tiling SARS-CoV-2 genome, resulting in a genome-wide distribution of well-defined structures as a function of position along the RNA. This distribution can be represented as a heatmap of well-defined structures along the full-length SARS-CoV-2 genome (Figure 3a), with expanded views of the initial two-thirds (5’UTR and ORF1ab, Figure 3b) and the downstream one-third of the RNA (Structural/accessory ORFs and 3’UTR, Figure 3c). Varying degrees of structural content are observed across the genome, ranging from 24% to 71% of well-defined structures in individual domains (Figure 4). To assess how the most highly structured regions are distributed throughout the genomic RNA, we started by analyzing the noncoding regions. As expected, we found a high level of structural content in each of the untranslated regions (the 5’ and 3’ UTRs): indeed, 61% of the 5’UTR and 41% of the 3’ UTR are characterized by well-defined structures (Figure 3a; Figure 4), which is consistent with the presence of RNA regulatory elements that play roles in replication and translation of the virus.

**Figure 3.**
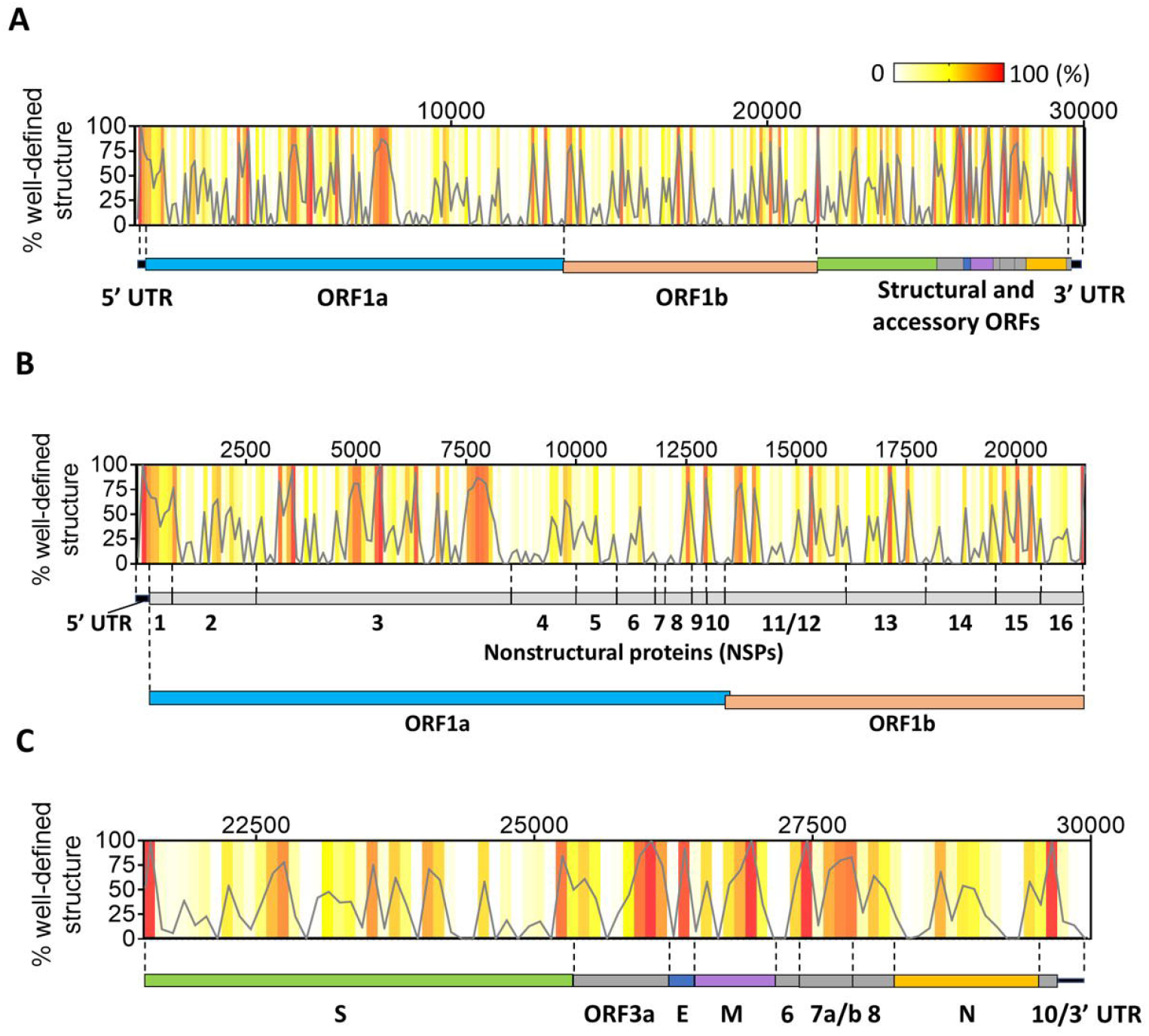
Distribution of well-defined RNA structures across the SARS-CoV-2 genome. (A) The percentage of nucleotides in well-defined structured regions (high BPC/low Shannon) was calculated in 100-nt bins tiling the genome and is plotted as a function of the genomic coordinate (gray curve). Individual percentages of each genomic bin are also represented as a heatmap in the same graph (color legend on the top right-hand corner). A scheme representing the genomic divisions of SARS-CoV-2 is shown next to the plot to guide location of structured regions. (B) An expanded view of the initial two-thirds of the genome from the graph in (A) is shown along with the genomic divisions of this region (UTR + ORF1ab and corresponding NSP divisions). (C) The downstream third of the genome is expanded from the graph in (A) to zoom in on individual structural and accessory ORFs in this region.

**Figure 4.**
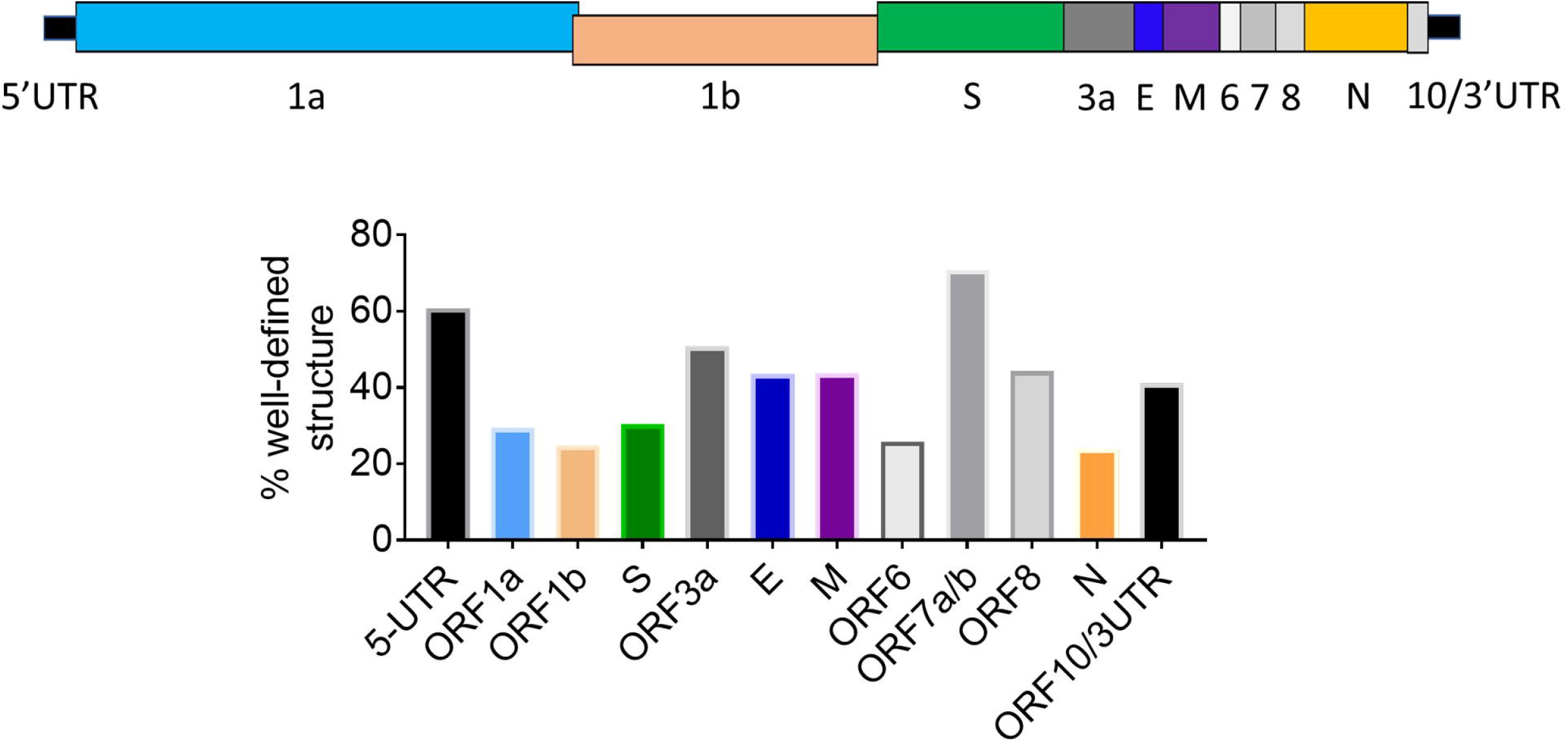
The percentages of nucleotides with well-defined structure (high BPC/low Shannon) are shown for each genomic section of SARS-CoV-2. A cartoon of genomic regions is depicted above the bar plot and each region is color coded relative to the bar graph.

However, unlike conventional mRNAs or flaviviral RNAs, stable RNA structural elements are not confined to the UTRs, as they are also observed in all coding regions of SARS-CoV-2 (Figure 3a). Different regions of the ORF contain varying amounts of stable secondary structure. For example, we observe that 27% of ORF 1ab contains nucleotides within high BPC/low Shannon regions, which are spread sparsely over more than 21 kb (~2/3 of the genome). These foci of well-defined RNA structures are not uniformly distributed along this ORF, as individual NSP domains contain different degrees of secondary structure (Figure 3b; Figure 5). The most upstream segment, NSP1, is also the most highly structured region of ORF1ab, with 56% of its nucleotides forming well-defined structures (Figure 5). Importantly, the upstream half of the NSP1 segment appears to be part of a large module that forms in conjunction with the 5’ UTR, as a peak of high BPC/low Shannon values that encompasses both domains (Figure 3b), suggesting that, as observed in other coronaviruses (Chen and Olsthoorn 2010), upstream regulatory elements of the genome extend far into the ORF. The largest domain in ORF1ab, NSP3, contains highly structured foci that can be organized into three big clusters (Figure 3b): one cluster located adjacent to the 5’ terminus of the domain (nt 3200-3600), a middle section that displays multiple high BPC/low Shannon peaks (nt 4500-6500) and a downstream segment near the 3’ NSP3 terminus (nt 7450-8200). This overall organization of NSP3 suggests the formation of independent modules of RNA secondary structure. Similar clusters of RNA structures are observed in NSPs 12 and 13 and they occur roughly within the limits of each domain. By contrast, other NSP regions (4, 5, 6, 8, 9, 10, 14, 15 and 16) form structures that encompass the boundaries of individual segments, suggesting a modular organization at the RNA level that does not necessarily correlate with functionality at the protein level. Finally, we observe that regions corresponding to NSP7 and NSP11 show a complete absence of well-determined structures (Figure 5), thereby demonstrating the presence of predominantly unstructured regions in the genome.

**Figure 5.**
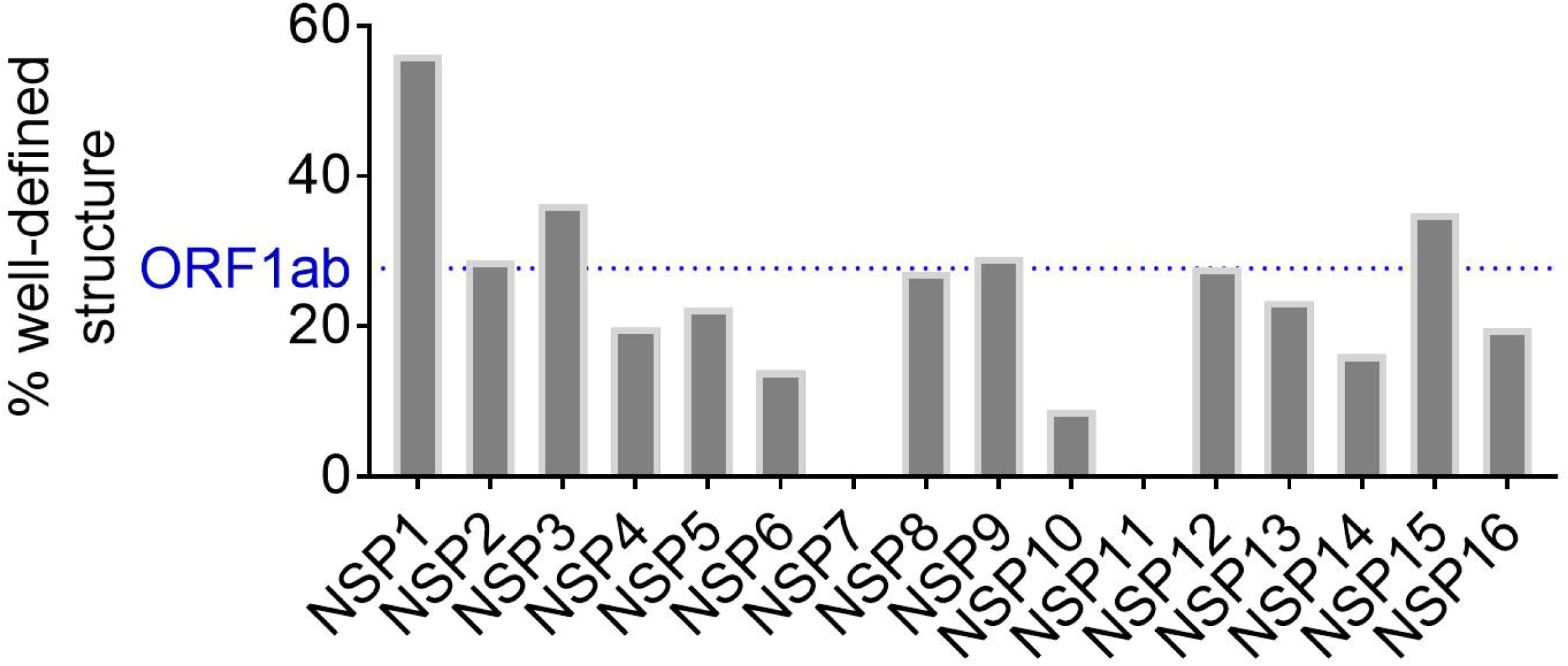
The percentages of nucleotides with well-defined structure (high BPC/low Shannon) are shown (gray bars) for each NSP (non-structural protein) section of SARS-CoV-2 ORF1ab. The horizontal dashed line (blue) represents the percentage corresponding to the entire ORF1ab.

The downstream third of the SARS-CoV-2 genome, which contains ORFs for structural and accessory proteins (the sgRNA-encoding region), displays a much higher overall secondary structural content than ORF 1ab, with a global fraction of well-determined structures of 36% (Figure 3c). Remarkably, some of these ORFs (3a, E, M, 7ab and 8) have a predicted structural content that is comparable or even higher than that of the UTRs (Figure 4), with the most prominent example being ORF7ab (high BPC/low Shannon fraction of 70%). These highly structured ORFs are all relatively short (ranging from 236 to 852 nt), consisting of a series of well-defined structures that are very closely spaced (Figure 3c). On the other hand, longer ORFs like S (Spike) and N (Nucleocapsid) contain shorter patches of well-defined structures interspersed with longer, less structured regions, resulting in a somewhat lower structural content for these ORFs (30% for the S ORF and 24% for the N ORF, Figure 4).

Similar to patterns observed for ORF1ab, we also observe RNA structural modules that span multiple ORFs. One example is a module that spans the junction between S ORF and ORF3a, including the TRS sequence at the intersection between them. Similarly, part of ORF 6 folds into a substructure that includes elements of ORFs 7a/b and 8, resulting in an extended structured region that includes three TRS elements. These observations indicate that some TRS sequences in this region are involved in structures with their surrounding ORFs, a feature that is likely to influence sgRNA synthesis and replication. Taken together, these results suggest the existence of numerous modules of well-determined RNA structure throughout the SARS-CoV-2 genome and they reveal important trends across functional domains.

### Secondary structural features depend on genomic vs subgenomic context

Intrigued by the abundance of predicted RNA structures within the ORFs of SARS-CoV-2, we asked whether individual ORFs might adopt different structures depending on the context in which they are inserted, i.e. in genomic vs subgenomic RNAs. As a direct application of base pair content analysis, we examined the predicted folding of one specific structural ORF that is present in both genomic and subgenomic RNAs. We chose the Nucleocapsid ORF because it forms the most abundant sgRNA (N sgRNA), which is estimated to be at least one order of magnitude more abundant than other sgRNAs (Kim et al. 2020).

When comparing the N ORF base pair content in both the genomic and subgenomic contexts (Figure 6a), one observes subtle differences in the upstream segment of this ORF. Specifically, the upstream 434 nucleotides of N ORF show patches of significantly higher base pair content in the subgenomic context than in the genomic context. To understand this, we examined the actual secondary structure predictions of this region in both contexts (Figure 6b). In the genomic context, sequences upstream of the N region (which belong to ORF8) are predicted to form base pairing interactions with the adjacent N ORF, resulting in a specific RNA structure that is uniquely dependent on the genomic environment. By contrast in the subgenomic context, upstream regions of N ORF are adjacent to the 5’ leader sequence, which folds somewhat autonomously into an independent motif and makes fewer contacts with the adjacent N sequences. As a result, upstream nucleotides of the N ORF form a compact alternative secondary structure in the sgRNA that is distinct, and which has a higher overall BPC than the same sequence in the genomic context.

**Figure 6.**
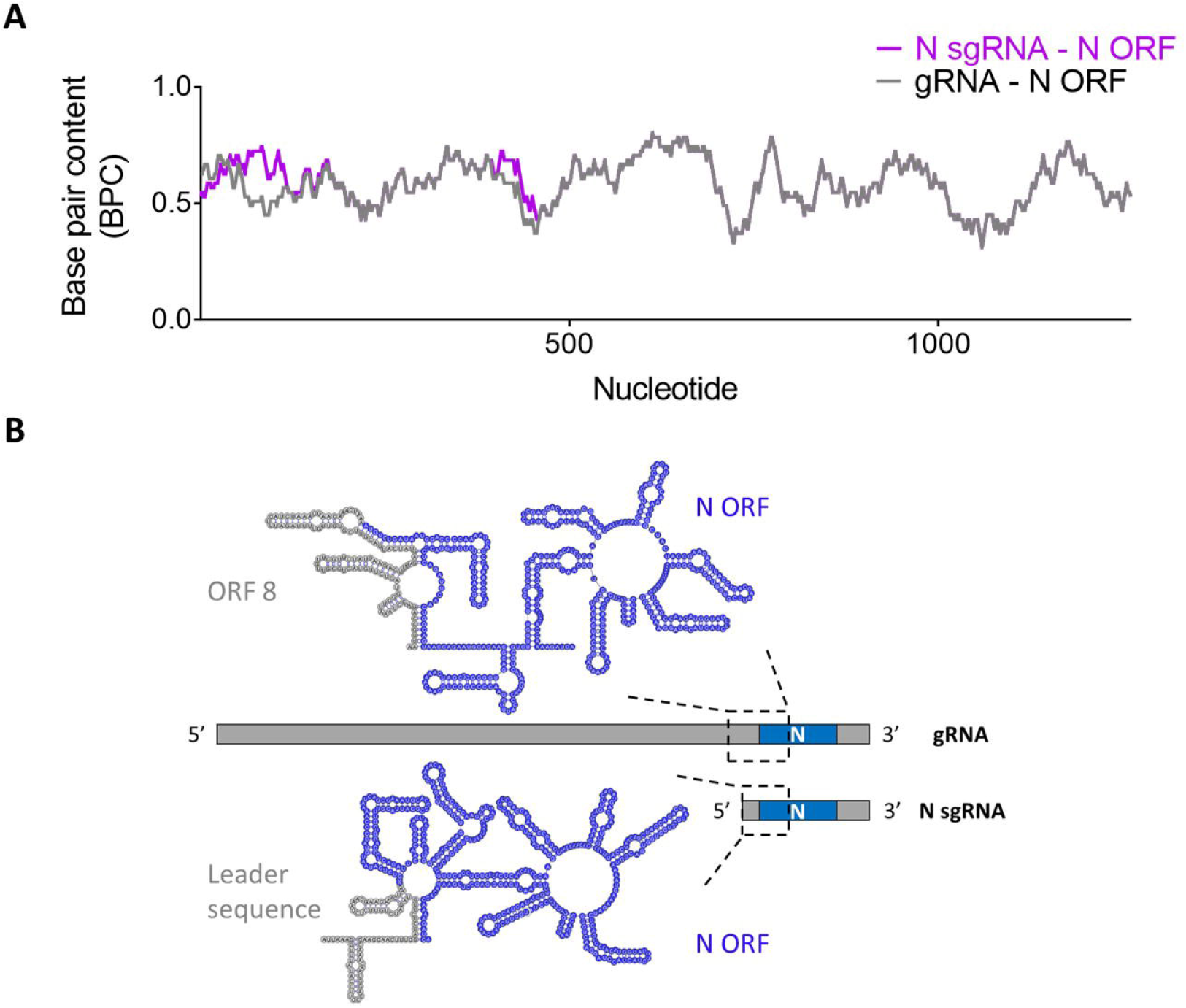
BPC analysis of the N ORF in genomic and N subgenomic contexts. (A) The base pair content for the N ORF (total of 1260 nucleotides) is plotted as a function of the nucleotide number for the genomic RNA (gray curve) and the N sgRNA (magenta curve). The x-axis numbering represents the N ORF nucleotide order (1-1260). (B) Secondary structure predictions containing the upstream 434 nucleotides of the N ORF are shown for both genomic and N subgenomic RNAs. The figure shows the region containing structural differences identified in (A). In the genomic RNA, the gray region represents a downstream segment of ORF8 and a stretch of additional sequence containing the TRS. In the N subgenomic RNA, the gray region is the 5’ leader sequence and a stretch of additional sequence containing the TRS. Structures were drawn on VARNA (Darty et al. 2009).

These results demonstrate that the same RNA sequences can adopt completely different structures in the subgenomic and genomic contexts, thereby diversifying functionality of the viral genome with significant implications for RNA stability, processing and molecular mechanism. A compact structure with high BPC in the upstream segment of the N sgRNA might contribute to its stability as an independent transcript, potentially explaining the unusually high abundance of this sgRNA. It will be important to conduct genetic studies to assess this model and to perform analogous studies on the other sgRNA/gRNA combinations to evaluate how they might influence viral function. These limited studies exemplify the presence of context-dependent differences in the structure of specific viral RNA sequences and they provide a framework for sifting through vast coronavirus genomes and identifying discrete elements of dynamic RNA secondary structure.

## Discussion

The majority of biophysical and structural studies being conducted on coronaviruses focus on the viral proteins, such as the virion constituents and components of the replication-transcription machinery (Gao et al. 2020; Wan et al. 2020; Wrapp et al. 2020; Zhang et al. 2020). However, RNA motifs within positive-strand RNA viruses guide many processes that are critical for the virus life cycle (You et al. 2004; McMullan et al. 2007; Friebe and Bartenschlager 2009; Modrow et al. 2013; Pirakitikulr et al. 2016). SARS-CoV-2 is unlikely to be an exception to this rule, particularly given that, to our knowledge, it has the most elaborately structured RNA genome that has ever been reported to date, and many of its constituent structures are conserved across coronavirus families (Chen and Olsthoorn 2010; Yang and Leibowitz 2015; Madhugiri et al. 2018; Rangan et al. 2020). Until the present study was conducted, the genome of HCV was the landmark example of a highly structured viral RNA, and one of the most highly structured ORFs in nature, distinguished by networks of functionally essential RNA structural motifs throughout both the coding and noncoding regions (Mauger et al. 2015; Pirakitikulr et al. 2016; Adams et al. 2017). However, we report here that the SARS-CoV-2 genomic RNA is nearly twice as compact and structured as HCV based on its folding stability, even when adjusting for its vastly greater overall length (~ 30 kb vs. ~10 kb). RNA structural motifs within the UTRs and ORFs of coronaviruses are seemingly larger and more complex than those observed in other virus families (Clyde and Harris 2006; Lim and Brown 2017; Li et al. 2018; Simon et al. 2019), suggesting that an understanding of coronavirus RNA structure will play a key role in understanding the mechanistic processes and vulnerabilities of this virus.

It is interesting to consider why the largest RNA genome might also be the most highly structured. One hypothesis is that the idiosyncratic SARS-CoV-2 genomic architecture serves a protective function. Biophysical studies have shown that extensively structured viral RNAs like HCV adopt highly condensed states in solution and that these are inaccessible to external probe hybridization (Davis et al. 2008). It is therefore reasonable to expect that the SARS-CoV-2 genomic RNA might adopt structural states that affect the way it interacts with both viral and cellular factors. The architectural features of the massive SARS-CoV-2 genome may confer protection against cellular nucleases, which would facilitate sustained infection in cells. In addition, the folded architecture of the SARS-CoV-2 genome may enable it to hide in plain sight, reducing activation of host pattern recognition receptors (Bowie and Unterholzner 2008). In other well-structured positive-stranded RNA viruses like HCV and MNV, the formation of genome-scale ordered RNA structure (GORS) correlates with decreased activation of antiviral pathways (McFadden et al. 2013; Tuplin 2015). Cell-based assays have shown that highly structured viral transcripts have a reduced propensity to activate interferon responses when compared with less structured viral RNAs (Witteveldt et al. 2014). Understanding the interplay between the SARS-CoV-2 genome architecture and elements of the host immune system will undoubtedly be a rich area of future investigation.

Another consequence of the highly structured, compact SARS-CoV-2 RNA genome is that its reduced dimensionality would facilitate interactions between RNA structural elements that are otherwise far apart from one another in primary sequence. Bringing genomic elements into close spatial proximity of one another will support the formation of long-range interactions between distant segments of the genome. Much like the topologically associating domains (TADs) in the chromatin of eukaryotes (Szabo et al. 2019), RNA TADs in viral genomes would have the capacity to control replication, translation, packaging, and many other processes, as suggested for structures that constrain ends of the HCV genome (Shetty et al. 2013; Romero-Lopez et al. 2014). There is already precedent for this within the coronavirus family, as long-range contacts in the TGEV genome and cross-talk between the 5’ and 3’ ends of the MHV genome have been proposed to modulate aspects of sgRNA synthesis in each system (Li et al. 2008; Sola et al. 2011).

Given the many mechanistic implications for “structuredness” of the SARS-CoV-2 RNA genome, we were motivated to adapt and develop tools for quantifying overall base pair content, and motif stability, relative to the expanse of an entire genome. For example, to monitor the domain-level distribution of extensively base paired regions across this RNA, we developed a general strategy for extracting the base pair content from a predicted secondary structure model using an approach that is readily applicable to any theoretical or experimentally-determined RNA structure prediction. By further applying a Shannon entropy filter (Mathews 2004), we were then able to focus our analysis on the regions of highest base pairing propensity and well-determined secondary structural composition (high BPC/low Shannon). Downstream quantification of the density of high BPC/low Shannon nucleotides enabled us to generate a profile of their distribution along the entire RNA. In this way, we could rapidly map regions with high and low RNA structural frequency along the SARS-CoV-2 genome, producing a snapshot of the structural landscape for an RNA of any size and pinpointing areas that merit focused empirical study. In massive RNAs, such as coronaviral genomes or certain eukaryotic mRNA transcripts, an approach that rapidly sifts information on structural content and puts it into a global and spatial context is vital for the discovery of regulatory modules and drug targets.

It is useful to reflect on the frequency and spatial distribution of secondary structures within SARS-CoV-2 genome, as their placement along the genome is far from uniform. Our analysis predicts that ORFs in the downstream third of the genome contain the highest density of well-defined structures in the viral transcript. These ORFs encode the sgRNAs and the accessory and structural proteins that are required during later stages of replication (Fehr and Perlman 2015; Kim et al. 2020). One possibility is that more extensive RNA folding of downstream segments might increase their relative stability and safeguard them for later phases of viral infection. Importantly, several RNA structures in this segment encompass the transcription regulatory sequences (TRS), which are key to production of the 3’-nested sgRNAs (Zuniga et al. 2004). Given that TRS elements mediate the fusion of each ORF terminus to the leader sequence during subgenomic replication, their structural context is likely to affect the frequency of template switching (leader-to-body fusion) at each fusion site, possibly involving interactions with the replicase-transcriptase complex or other gRNA-interacting partners like the nucleocapsid protein (Hurst et al. 2013; McBride et al. 2014). The numerous structures found within the coding sequences of this region may also contribute to processes other than RNA synthesis, such as translational regulation (Clyde and Harris 2006; Jaafar and Kieft 2019) and infectivity (Pirakitikulr et al. 2016).

It is notable that some of the structures predicted in downstream regions of the genome will depend on transcript context, as many of them occur at junctions between consecutive ORFs. Many potential structures will no longer form after leader-to-body fusion occurs at the junctional TRS sites (upon formation of an sgRNA), suggesting that certain structures may have roles only in the context of full-length genomic RNA and/or in longer sgRNAs that arise from upstream fusion events. We explore one example of such a structure (Figure 6), which involves base pairings between a downstream segment of ORF 8 and upstream segments of the N protein ORF, which are ablated upon formation of the N sgRNA. If functional, genomic structures of this type may facilitate processes such as viral packaging (Masters 2019) and they may promote infectivity (Smyth et al. 2018). On the other side of the spectrum, secondary structures that form exclusively in sgRNAs, such as the large motif predicted between the 5’ leader sequence and the N ORF (Figure 6b), are expected to affect sgRNA properties like stability, abundance, and the recruitment of sgRNA-specific factors. Systematic structural comparisons among SARS-CoV-2 transcripts will certainly help identify candidate genomic structures with potential roles in infectivity and to provide a framework for rationalizing the relative stabilities and functions of sgRNAs.

The vast genome of SARS-CoV-2 and its complex transcriptome present new challenges to RNA science, immunology and medicine. However, the SARS-CoV-2 system and the intense attention it has attracted will also stimulate innovation, pushing researchers to develop new strategies for addressing the many challenges of studying and understanding exceptionally large RNA transcripts, particularly those in the lifecycle of pathogens. We hope that the results and methods described in this work will facilitate the design of new experiments for understanding the modular architecture of the SARS-CoV-2 genome and for unraveling the complex mechanisms of viral pathogenicity and host response. In addition, by focusing on the most structured regions of the genome and characterizing their constituent motifs, we hope to stimulate the search for promising new drug targets, leading to novel therapeutic strategies against COVID19.

## Methods

### Minimum free-energy (MFE) Z-score analysis

Folding minimum free-energy (MFE) Z-scores for SARS-CoV-2, HCV, WNV and human mRNAs (GAPDH, ACTB, HPRT and a-tubulin) were calculated with the ScanFold-Scan program (Andrews et al. 2018). ScanFold-Scan uses the ViennaRNA Package 2.0 (Lorenz et al. 2011) to fold the target sequence in sliding windows and calculate the MFE secondary structure for each window. The Z-score for each window is computed by calculating by the difference between the native MFE and the average MFE of shuffled sequence controls and normalizing this difference by the standard deviation of the shuffled MFE distribution. ScanFold-Scan default parameters were used (120 nt window size, folding temperature 37°C, mononucleotide shuffle procedure), with the exception of the number of randomizations (set to 100) and the sliding step size (set to 1nt). Z-score frequency distributions for SARS-CoV-2, HCV, West Nile Virus and the composite of human mRNAs were calculated on GraphPad Prism.

### RNA secondary structure modeling

Secondary structure predictions for full-length SARS-CoV-2 RNA genome and the Nucleocapsid (N) subgenomic RNA were obtained using SuperFold with default settings as described (Smola et al. 2015). Since no experimental constraints were used in the modeling, the SHAPE contribution was canceled by setting both the SHAPE slope and intercept to 0 (--SHAPEslope 0; --SHAPEintercept 0). Briefly, SuperFold calculates the base pairing partition function in 1200 nt windows in steps of 100 nt along the RNA, while removing interactions occurring within the terminal 300 nt at the 5’ and 3’ ends of each window; to compensate for the de-weighting at the true 5’ and 3’ ends of the RNA, additional partition function calculations on those termini are performed. Base pairing probabilities are averaged across all windows in which a base pair is predicted to form. SuperFold then uses the Fold function from RNAstructure (Reuter and Mathews 2010) to calculate the minimum free-energy secondary structure in 3000 nt sliding windows every 300 nt along the RNA; highly probable base pairs from the partition function calculation (P>99%) are used as hard constraints in this step. The default maximum base pairing distance (600 nt) was used in both partition function and windowed folding steps. Finally, a consensus secondary structure is obtained by outputting base pairs consistently predicted during windowed folding, i.e. occurring in more than one-half of windows. The Shannon entropy for each nucleotide is computed with base pairing probabilities derived from the partition function calculation.

### Base pair content (BPC) and relative base pair content (BPC_rel_) calculations

The base pair content (BPC) was calculated from the predicted secondary structure in sliding windows of 51 nt tiling the RNA in steps of 1 nt. For each nucleotide, we define BPC as the fraction of base paired nucleotides within the window centered about that nucleotide. For the terminal 25 nucleotides at both ends of the RNA (window sizes < 51 nt), sliding windows were truncated accordingly.

The BPC_rel_, i.e. the relative base pair content for a given nucleotide, was calculated by computing the percentile of the nucleotide’s absolute BPC among the global set of BPC values comprising the entire RNA. Briefly, 100,000 values were resampled with replacement (bootstrapping) from the set of BPC values. The bootstrapping is used as a convention to approximate the percentile of BPC values across a common denominator, regardless of the

RNA tested. For each nucleotide, we then computed the fraction of bootstrapped values that lie below that nucleotide’s BPC score. By doing this, the median BPC value for a given RNA is “normalized” to 0.5, making for an intuitive measure of the relative significance of the magnitude of a given nucleotide’s BPC value. In case the median value itself is present multiple times in the BPC dataset, the standardized median threshold shifts beyond 0.5, which can be corrected by resetting all median BPC values with a BPC_rel_ of 0.5. In this way, the rescaled BPC values (BPC_rel_) can be used for direct quantitative comparison of the structural content across any RNA.

### Identification of well-defined structures (high BPC/low Shannon)

Base pair content (BPC) along with Shannon entropy values were used to identify nucleotides likely to form well-defined structures. In SHAPE experiments, this is accomplished by flagging those nucleotides with both low SHAPE reactivity and low Shannon entropy values, after smoothing both datasets in sliding windows tiling the RNA in steps of 1 nucleotide (Smola et al. 2015; Smola et al. 2016). Nucleotides with SHAPE reactivities and Shannon entropy values below the global median of each respective distribution are considered highly structured and well-defined. By analogy with the “low SHAPE/low Shannon” concept, here we define “high BPC/low Shannon” nucleotides as those likely engaged in well-defined structures. Shannon entropy values were first smoothed in 51 nt sliding windows (steps of 1 nt) to match the BPC calculation parameters. After computing relative BPC values (BPC_rel_), nucleotides with BPC_rel_ greater than 0.5, i.e. above the global median of the distribution, and Shannon entropy values below the global median were flagged as “high BPC/low Shannon”. The fraction of well-defined structure in a given section of the RNA was then defined as the fraction of “high BPC/low Shannon” nucleotides in that section.

### Overlap between high BPC/low Shannon regions and regions with low average Z-scores

It was important to assess the overlap between nucleotides in well-defined regions as defined by our approach (high BPC/low Shannon) and nucleotides with low average Z-scores (Z_avg_) from the ScanFold-Fold analysis of SARS-CoV-2 reported in Andrews et al. (Andrews et al. 2020). To this end, Z_avg_ scores for SARS-CoV-2 were downloaded from the Moss lab RNAStructuromeDB (https://structurome.bb.iastate.edu/). In order to match the criteria we used to flag well-defined structures (high BPC/low Shannon, both relative to the global median), low Z_avg_ values are defined here as those occurring below the distribution median of Z_avg_ values, i.e. regions with folding stability above the average. The statistical significance of the overlap between both methods was evaluated on Matlab by running simulations of randomly distributed elements in both groups. The P-value was estimated by computing the number of times an overlap equal or greater than the observed value was obtained and then dividing it by the number of simulations.

### Reference sequences

The SARS-CoV-2 reference genome from Wu et al. (Wu et al. 2020) was used for all analyses, along with the protein annotations deposited in NCBI (accession IDs MN908947.3 and NC045512.2). Human beta-actin mRNA (NM_001101.5), human GAPDH mRNA (NM_002046.7), human HPRT mRNA (NM_000194.3), human α-tubulin 1 mRNA (NM_006009.4), West Nile genome (NY99 sequence, based on DQ211652.1 reference) and HCV genome (JC1 sequence, based on JF343782.1 reference) sequences were used for Z-score analysis.

### Data availability

Data sheets, sequence files and scripts used in this work are available at the GitHub repository: https://github.com/pylelab/SARS-CoV-2_global_local_structure

## Acknowledgments

We would like to thank Han Wan (Pyle lab, Yale University) for thoughtful comments on the manuscript. This work was supported by the Howard Hughes Medical Institute (A.M.P.) and a Yale College Dean’s Research Fellowship (to G.M.). Funding for open access charge was provided by the Howard Hughes Medical Institute.

